# Entorhinal neurons exhibit cue locking in rodent VR

**DOI:** 10.1101/477620

**Authors:** Giulio Casali, Sarah Shipley, Charlie Dowell, Robin Hayman, Caswell Barry

## Abstract

The regular firing pattern exhibited by medial entorhinal (mEC) grid cells of locomoting rodents is hypothesized to provide spatial metric information relevant for navigation. The development of virtual reality (VR) for head-fixed mice confers a number of experimental advantages and has become increasingly popular as a method for investigating spatially-selective cells. Recent experiments using 1D VR linear tracks have shown that some mEC cells have multiple fields in virtual space, analogous to grid cells on real linear tracks. We recorded from the mEC as mice traversed virtual tracks featuring regularly spaced repetitive cues and identified a population of cells with multiple firing fields, resembling the regular firing of grid cells. However, further analyses indicated that many of these were not, in fact, grid cells because: 1) When recorded in the open field they did not display discrete firing fields with six-fold symmetry; 2) In different VR environments their firing fields were found to match the spatial frequency of repetitive environmental cues. In contrast, cells identified as grid cells based on their open field firing patterns did not exhibit cue locking. In light of these results we highlight the importance of controlling the periodicity of the visual cues in VR and the necessity of identifying grid cells from real open field environments in order to correctly characterise spatially modulated neurons in VR experiments.

## Introduction

Since their discovery, the striking regularity of grid cell firing patterns has been proposed to play a role in encoding travelled distances and are widely held to be a core component of a circuit necessary for the integration of self-motion cues – ‘path integration’ (Burak and Fiete, 2009; Burgess, 2008; Hafting et al., 2005; McNaughton et al., 2006; Winter et al., 2015). Equally, the influence of the sensory environment on grid cell firing is also well established. In rodents, manipulations made to familiar spatial cues result in commensurate changes to grid-patterns (Hafting et al., 2005; Barry et al., 2007; Stensola et al., 2012), in geometrically polarised environments firing is distorted (Krupic et al., 2015; Stensola et al., 2015; Krupic et al., 2018), and in darkness their spatially periodic activity can break down completely (Chen et al., 2016; Pérez-Escobar et al., 2016).

Hence, it appears that while grid cell activity is shaped by self-motion information (Winter et al., 2015), sensory access to landmarks is necessary to maintain stable spatial firing (Campbell et al., 2018; Hardcastle et al., 2015; Muessig et al., 2015). Further, the relative efficacy of these two sources of information (‘self-motion’ vs ‘landmark’) appears to change dynamically with experience. For example, when rats first explored a pair of visually identical enclosures connected by a corridor, grid cell firing in the enclosures was highly similar, suggesting a dominance of landmark-based information. However, with prolonged experience, grid-patterns disambiguated the two sides, forming a single global representation of the space, consistent with increasing emphasis being placed on self-motion cues (Carpenter et al., 2015). Similarly, computational work has also highlighted the necessity of landmark information as a means to reset accumulated errors in path integration networks (Banino et al., 2018; Burgess and Burgess, 2014). However, the mechanisms by which this reset occurs and by which the relative importance of different information sources can be modulated remains unclear.

Rodent virtual reality (VR) provides a powerful and increasingly popular experimental tool capable of manipulating the characteristics of an animal’s environment, thus offering an ideal means to address such issues (Thurley and Ayaz, 2017). Indeed, a number of studies have examined the responses of neurons recorded from the entorhinal and hippocampal networks in both 1D (Harvey et al., 2009; Dombeck et al., 2010; Domnisoru et al., 2013; Schmidt-Hieber and Häusser, 2013; Campbell et al., 2018) and 2D (Aronov and Tank, 2014; Chen et al., 2018). However, many cell types in these brain regions (e.g. grid cells, boundary vector cells and, to a certain extent, place cells) are typically identified from their open field firing patterns recorded during real environment foraging tasks. As such, it can be challenging to positively identify a neuron recorded solely from a 1D VR recording.

Therefore, to better understand how landmark and path integration information interact we recorded mEC grid cells as head-fixed mice ran through three distinct 1D VR environments, each consisting of different sets of regularly repetitive spatial cues. Our rationale was to explore the effects of repetitive spatial cues on grid and non-grid cell activity. Subsequently cells were classified as grid or non-grid cells based on their activity recorded during foraging in real 2D environments. In the case of grid cells, we found no evidence that grid-patterns were reset to a constant phase by each cue occurrence, thus pointing to a strong influence of path integration cues. However, we identified a population of non-grid neurons that did exhibit pronounced cue-locking, firing with a constant spatial relationship to each cue occurrence. In the repetitive environments used here, these cells appeared to have strongly periodic spatial firing and were erroneously categorised as grid cells by a measure previously used to identify grid cells under such circumstances (Domnisoru et al., 2013). We highlight the importance that 2D recordings play in the positive identification of spatial cell types that, like grid cells, are defined by their functional properties. Moreover, the presence of a class of neurons in mEC responding solely to visual landmark cues is novel and raises several questions regarding their contribution to the cognitive spatial network which we aim to answer with future experiments.

## Results

In total, 690 unique superficial mEC neurons were recorded from 4 male C57BL/6 mice during the course of 12 experimental sessions. At the end of each recording session tetrodes were advanced by a minimum of 50μm to avoid sampling of the same cells. *Post-hoc* examination of histology confirmed tetrode track location in superficial layers of left mEC.

In each session, mice initially foraged in a real world (RW) 2D environment for randomly thrown drops of soya milk (SMA, Wysoy), and after a minimum break of 10 minutes were head-fixed and left in darkness for about 5 minutes before a virtual reality (VR) session commenced (**Supp. Fig 6**).

In each session, animals ran in three different 1D VR environments (A, B, and C), performing traversals for liquid reward delivered at a fixed goal location near the end of the track before being teleported back to the start of the track. Each environment was comprised of repeated segments containing a number of visual cues (**Supp. Fig 2**). Trials in each environment were a minimum of 20 minutes duration.

In total 15 grid cells from 2 animals were identified based on spatial firing in the RW open field (gridness score vs. 95^th^ percentile of a shuffled distribution, mean grid score of identified cells = 0.58 ± 0.07, see Methods). Visual inspection confirmed that on the VR linear tracks these grid cells exhibited multiple spatially localised firing fields (**Figure 1A**) having similar firing rates between VR and RW (mean firing rate: RW = 1.3 ± 0.2 Hz, VR = 1.1 ± 0.4 Hz, Wilcoxon signed ranked test: *P* = 0.07; peak firing rate: RW = 5.4 ± 1.6 Hz, VR = 4.7 ± 1.5 Hz, Wilcoxon signed ranked test: *P* = 0.08) but reduced stability in VR compared to RW (first vs second half correlation: RW = 0.48 ± 0.05, VR = 0.07 ± 0.03, Wilcoxon signed ranked test: *P* = 0.0005).

**Figure 1.**
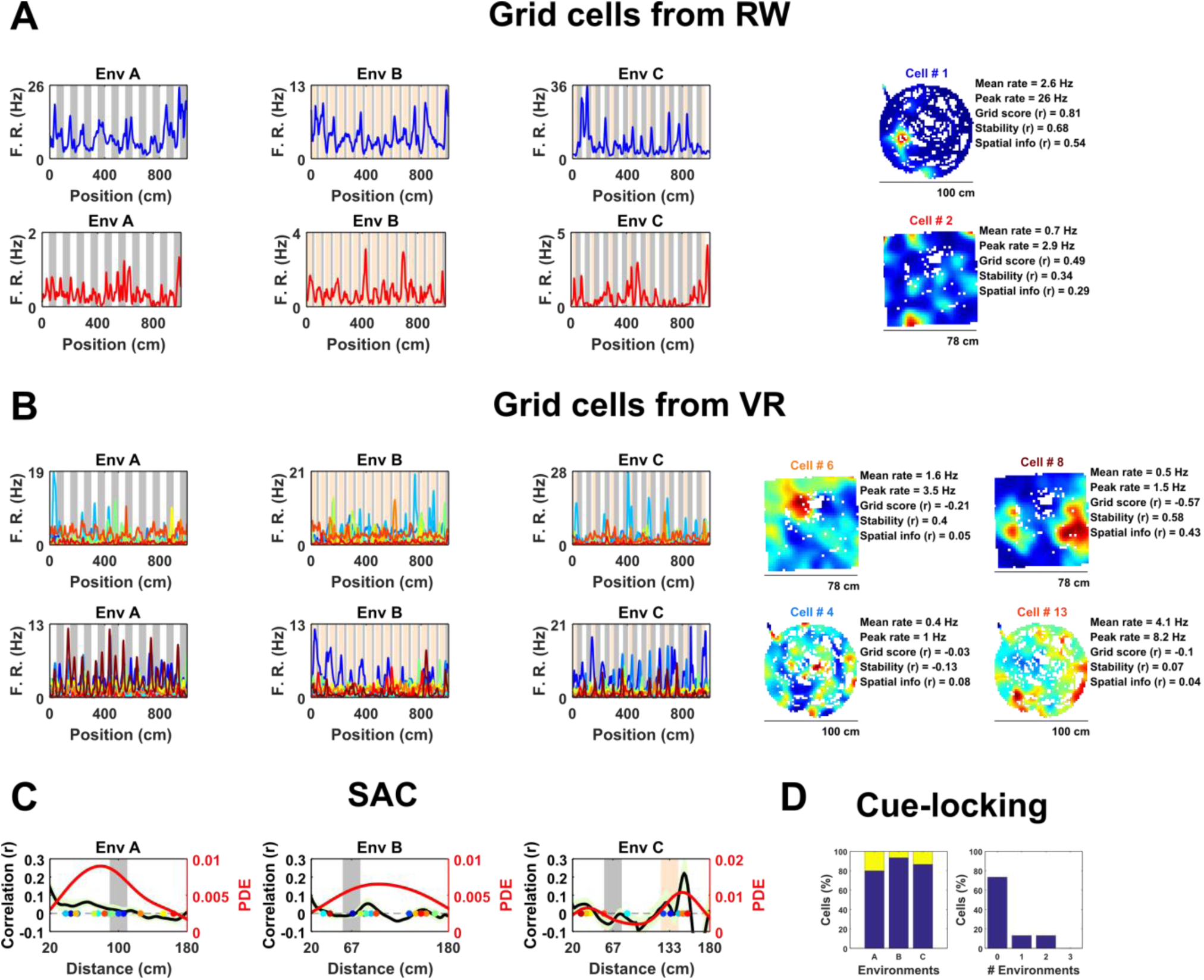
Spatially periodic activity in real (RW) and virtual (VR) environments, a subset of non-grid mEC cells show cue-locking in VR environments. **A,** Spatial activity from two example grid cells – identified based on RW recordings (*right*) – in each of the three VR environment (*left*). Repeating cues are indicated as coloured bars in the background of the VR plots. Cells are colour coded such that the title on RW ratemaps matches line colour on the VR plots. **B**, Similar to **A**, spatial activity from two cells which were (incorrectly) classified as grids cells based on VR activity (*left*) but not based on RW open field activity (*right*). Despite the regularity of their firing patterns in VR, these cells showed no clear grid-like firing in RW and only limited spatial responses. **C**, Cue-locking in grid cells (n=15, identified from RW) was investigated using spatial auto-correlograms (SACs). Plots show mean (black line) ± SEM (light green shade area) SACs across cells (left y-axis). Note the lack of periodicity corresponding to the frequency of cues in the VR environments (indicated by grey and orange bands). The overlaid color-coded dots represent the dominant spatial frequency in the 20-180cm range detected from the SAC of each grid cell – the distribution of these points is indicated by the red line (right y-axis). **D**, Proportion of grid cells exhibiting cue-locking (yellow) and no cue-locking (blue) in each VR environment (*left*) and proportion showing cue-locking in multiple VR environments (*right*).

Surprisingly, based on their activity in VR, relatively few of these cells were positively identified as grid cells. Specifically, using a method for grid cell categorisation developed for VR environments less repetitive than the ones used here (Domnisoru et al., 2013), in environment A, 2 of 15 cells were correctly classified as grid cells, in B 5 out of 15, and in C 3 of 15. These proportions which were not dissimilar to that of the whole ensemble of mEC cells: A = 106/690, binomial test: *P* = 0.99, B = 145/690, binomial test: *P* = 0.17, C = 108/690: binomial test: *P* = 0.45. Equally, the same criteria classified 74 of the 690 (10.7%) mEC neurons as grid cells based on their 1D VR activity, only 2 of which were also classified as grid cells based on their RW open field activity (gridness > 95^th^ of shuffle) (**Figure 1B**; grid score = −0.11 ± 0.03, stability = 0.17 ± 0.03, spatial information = 0.34 ± 0.05). Taken together, these numbers highlight the difficulty – and in particular high false positive rate – inherent in the identification of grid cell firing from 1D spatial data.

Next, we examined the extent to which grid cell activity was modulated by proximity to landmark cues in the VR. For each cell we calculated the mean spatial autocorrelogram (SAC) across trials in each VR environment and then detected the dominant spatial frequency - the highest peak in the 20-180 cm range (**Figure 1C**). The number of grid cells exhibiting ‘cue-locking’ in the VR environments – having a dominant spatial frequency matching that of the cues (see Methods) - was no greater than expected by chance (**Figure 1D**; A = 3/15, binomial test: *P* = 0.41; B = 1/15, binomial test: *P* = 0.14; C = 2/15, binomial test: *P* = 0.38). Indeed, 11 grid cells were not cue-locked in any of the VR environment, 2 were cue-locked in a single environment and 2 were cue-locked in 2 environments, with none of the grid cells being cue-locked in all three environments. Thus, grid cells – identified from the open field – showed no obvious tendency to be reset or modulated by the repetitive visual cues.

Next, we examined the spatial activity of all non-grid mEC neurons in VR (n = 675). To classify these neurons we developed a method analogous to a 1D version of the widely used gridness metric derived from the mean SAC (Sargolini et al., 2006). Briefly, cells with a maximum peak in the mean SAC exceeding the 95th percentile of single-cell shuffled distribution in at least two of the three VR environments were considered to exhibit significant spatially periodic activity. Our analysis detected 56 such cells, significantly more than expected by chance (chance = 5/675, binomial test, *P* < 0.0001). Inspection of the 1D ratemaps of these cells revealed pronounced and regular modulation of their spatial activity across VR environments, whereas their firing patterns in RW indicated that these neurons were unlikely to be un-detected grid cells (**Figure 2**, **Supp. Fig. 3**).

**Figure 2.**
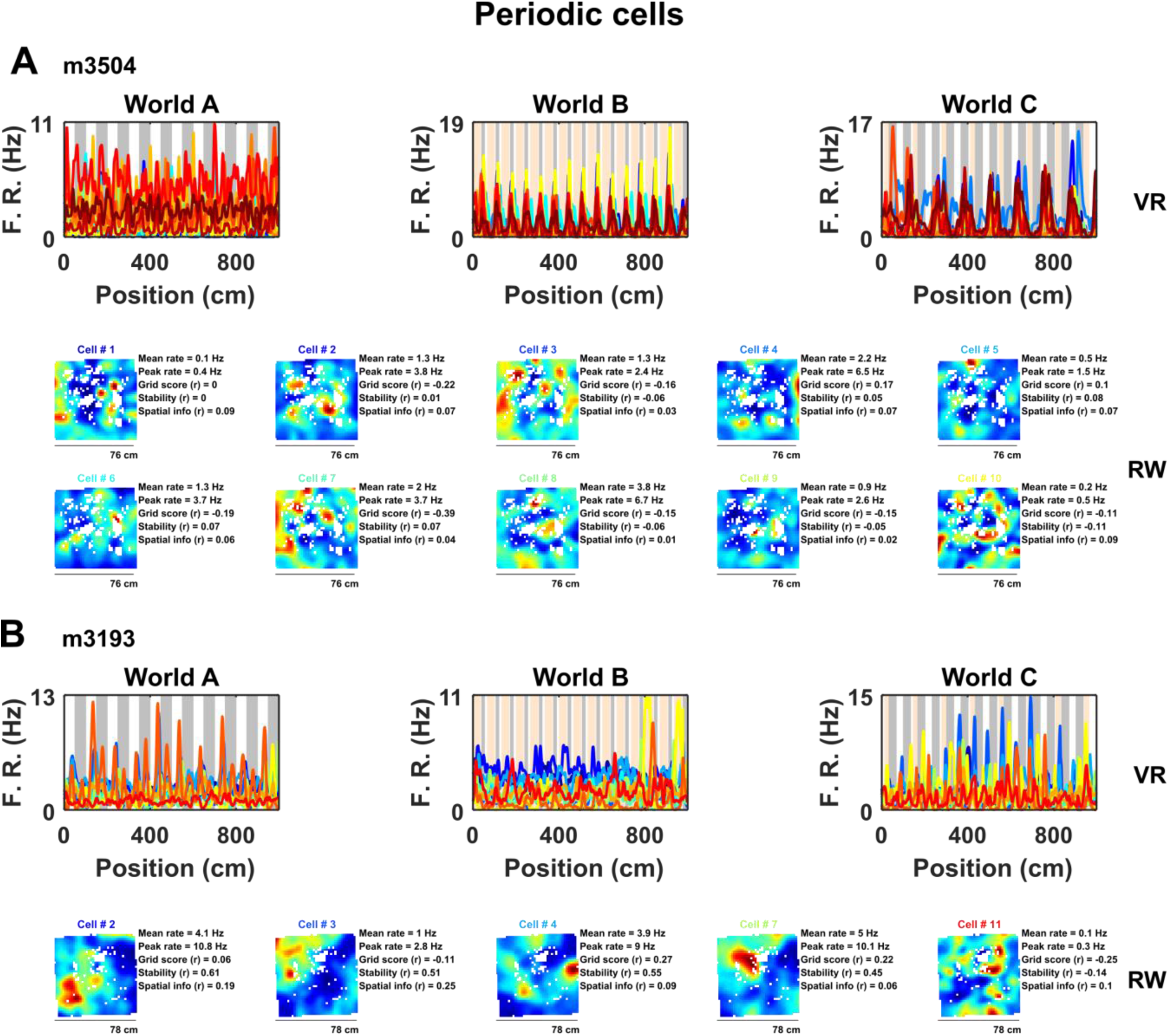
A sub-population of non-grid cells exhibit pronounced spatial periodicity in 1D VR environments. Examples of co-recorded non-grid periodic cells from 2 animals (**A**, 10 cells from mouse m3504: **B**, 5 cells from mouse 3193) shown as ratemaps in both VR (*top row*) and RW (*bottom rows*). In VR, the firing rate of cells is plotted against position in each environment (**A-C**) with the periodicity of repeating cues indicated in the background (grey and pink). In RW the ratemaps of the same color-coded cells in VR are shown together with mean and peak firing rate, grid score, stability, and spatial information. Despite the regularity of the firing pattern patterns in VR, these cells neither showed clear grid-like firing pattern nor spatial firing of any kind in RW.

Examination of their spatial firing in RW open field showed that these cells exhibited weakly spatial activity (0.22 ± 0.04 bits/spike) and no evidence of six-fold symmetry (0/56 cells exhibited significant grid scores vs. 95th percentile of a shuffled distribution, grid score = −0.13 ± 0.03, spatial stability = 0.14 ± 0.03, **Figure 2**,). However, a large portion of these cells (27/56 = 47%) passed criteria for grid cell inclusion based on previously used methods to detect grid cells on 1D VR linear tracks (Domnisoru et al., 2013).

To better characterize the nature of these non-grid periodic cells we focused on the relationship between the visual cues in VR and their regular firing patterns. We observed that within each VR environment the mean SAC was remarkably alike across cells, having a similar spatial frequency **Figure 3**). However, the periodicity of their activity differed between environments, and in most cases coincided with the underlying spatial frequency of the repetitive cues in those environments (**Figure 3A**). To quantify this observation, we repeated the analysis conducted to identify modulation of grid cells by visual cues - detecting the spatial frequency of each cell from the peak in the mean SAC and comparing that with the spatial frequency of cues in each environment (**Figure 3B**)– finding that these cells were strongly ‘cue-locked’ (proportion cue-locked cells: environment A = 28/56, binomial test: *P* < 0.0001; B = 34/56, *P* = < 0.0001; C = 43/56, *P* < 0.0001). Moreover, only 7/56 of the periodic cells were not cue-locked in at least one of the three environments, with 16/56 being cue-locked in 1 environment, 27/56 in 2 environments and 6/56 in all 3 environments (**Figure 3C**).

**Figure 3.**
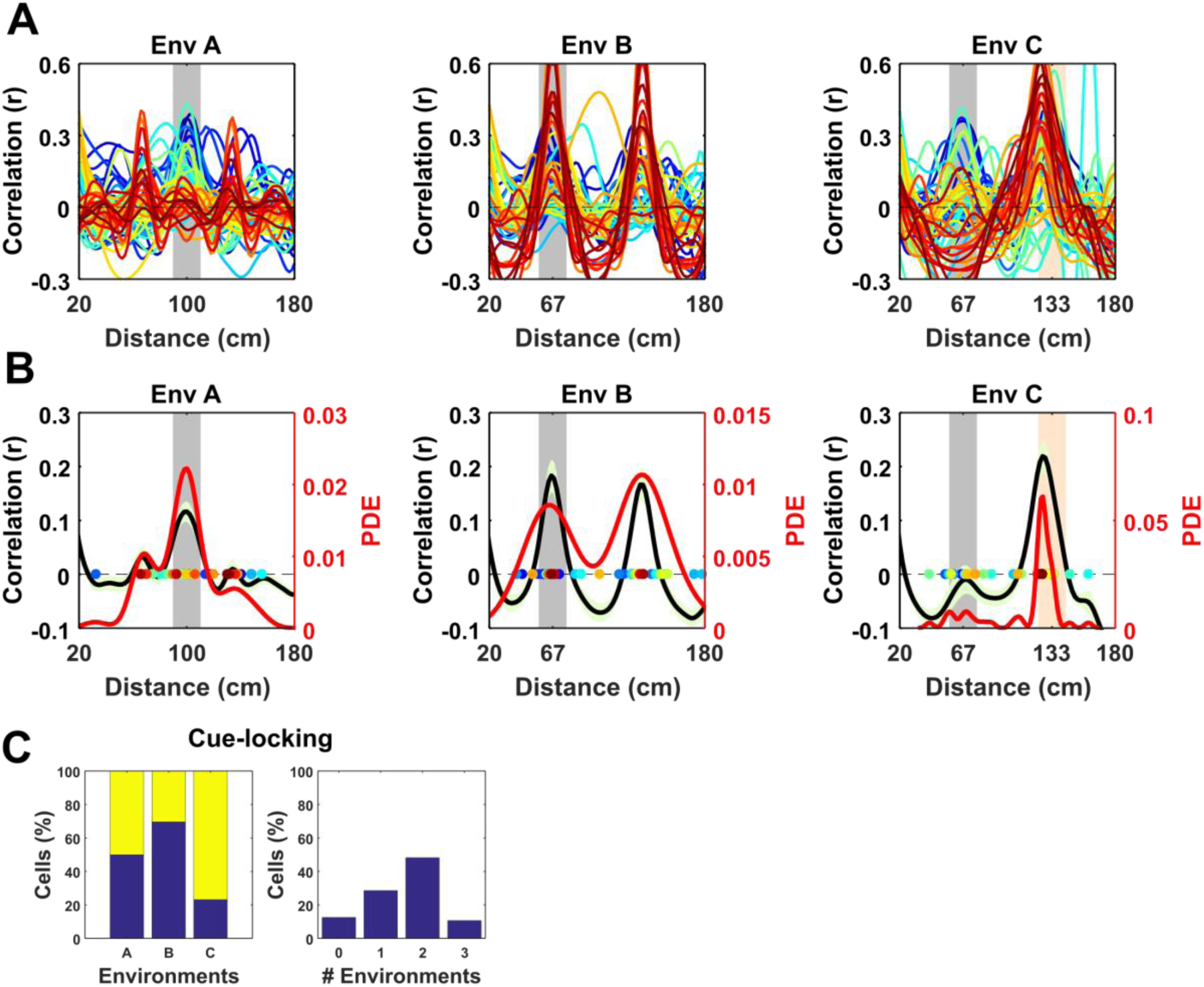
Spatial frequency of the periodic non-grid cells. Within VR environments, periodic non-grid cells exhibited regular firing at the same spatial frequency as the underlying repetitive visual cues, unlike grid cells which showed weaker spatial periodicity of varying frequencies. **A**, SAC of all non-grid periodic cells across VR environments. Note the clear peaks centred on the spatial frequency of the repeating cues of each environment (grey block). **B**, Mean (black line) ± S.E.M. (light green shaded area) of the SAC (left y-axis) across cells within each environment showing clear coincidence with the frequency of the repetitive cues (grey block). Coloured dots indicate the dominant spatial frequency of each cell (colour matches lines in **A**) and were used to compute the kernel density estimate (red line, right y-axis). **C**, Histograms showing (*left*) percentages of non-cue locked (blue) and cue-locked (yellow) periodic cells within each VR environment. *Right*, Histograms showing percentages of periodic cells exhibiting cue locking in multiple environments. Most cells (>85 %) displayed cue-locking in at least one VR environments.

Together, these results suggest that the regular firing pattern exhibited by these periodic cells was strongly modulated by the repeating visual cues rather than reflecting an internally-generated path integration signal like the one hypothesized for grid cells. In support of this notion, the proportion of periodic cells exhibiting cue-locking was significantly higher than the proportion of cue locked grid cells in environments B and C (χ^2^ = 10.62, *P* = 0.0011; χ^2^ = 20.5, *P* < 0.0001) though not A (χ^2^ = 3.71, *P* = 0.054).

Having identified a subset of non-grid cells exhibiting strong cue modulation, we next examined how their spatial responses were distributed relative to visual cues. Since the VR environments were composed of repeating linear segments, we calculated for each cell its mean rate map over the repeating unit. Visualised in this way it was clear that the spatial periodic firing of different cue locked cells had variable phases relative to the visual cues (**Figure 4A**). Moreover, when rate maps were sorted according to the location of their peak activity it was apparent that there was no strong tendency for firing to cluster at specific phases of the repeated segments (**Figure 4B;** Rayleigh test for peak density: VR environment A: *P* = 0.99, B: *P* = 0.99, C: *P* = 0.88). Finally, we examined if the relative phases at which non-periodic cells were active was conserved across environments. To test this, we focused on environments A and B which had the simplest patterns of repetitive visual cues, and cross-correlated the stacked rate maps from both environments sorted by peak location in A (**Figure 4C**). The resulting cross-correlation exhibited a single predominant peak (*r* = 0.25) which exceeded the values obtained by randomly shuffling the order of the cells before cross-correlating (*n* shuffles = 1000, peak from shuffle *r* = 0.014 ± 0.002, one-sample t-test, *t*_999_ = −101.1, *P* < 0.0001). Taken together these results indicate that although non-grid periodic cells are strongly modulated by environmental cues there is tendency for the relative phase at which cells fire to be preserved across environments.

**Figure 4.**
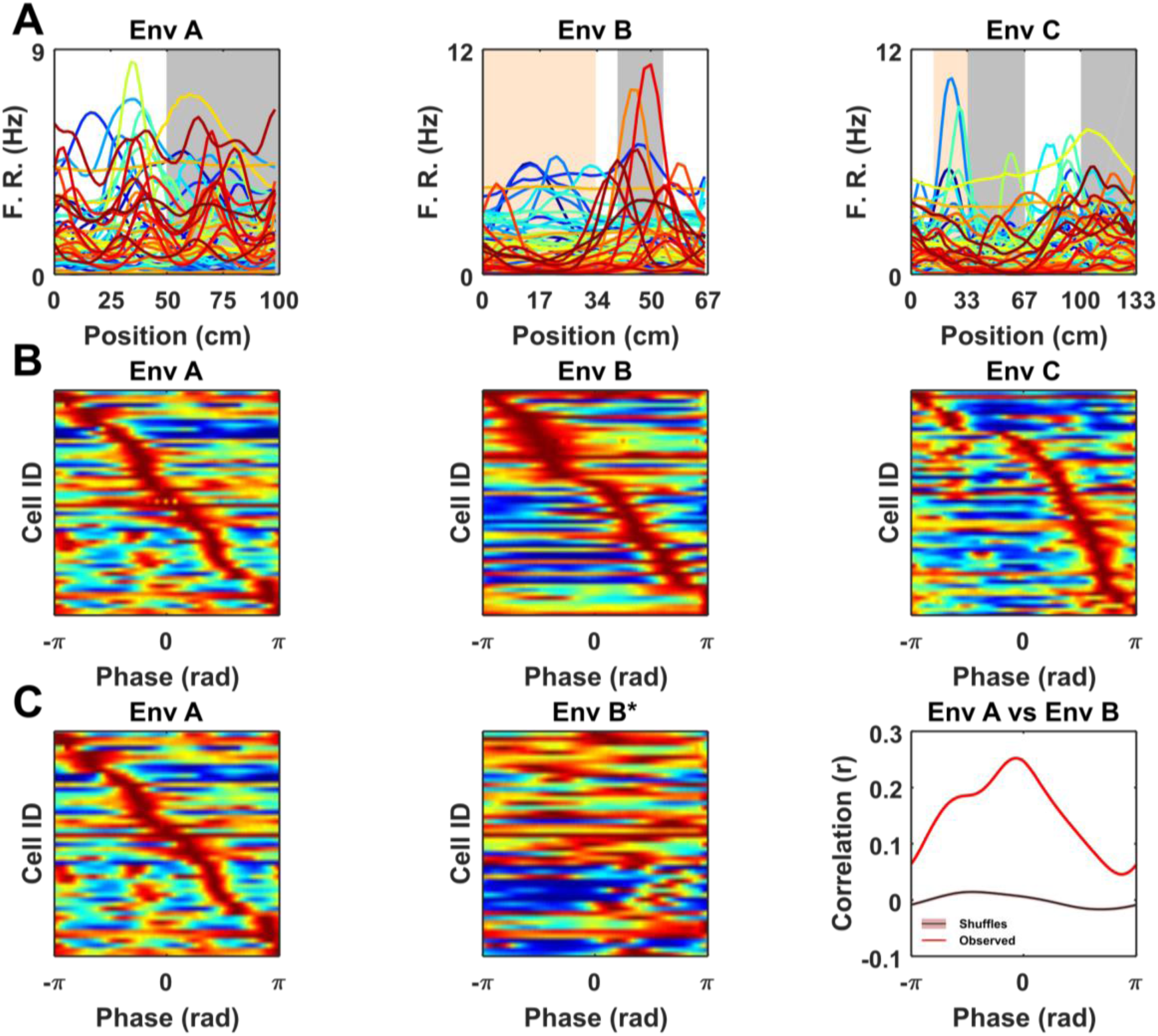
Non-grid periodic cells were strongly modulated by the frequency of the repetitive segments in each VR but were not clustered at specific phases within each segment. **A**, Rate maps for each color-coded cell across VR environments showing mean firing rate as function of location within the repeating segment. Note the differences in the peak firing rate and location of the spatial tuning curves across cells. **B**, Ratemaps of all cue-locked cells sorted according to location of peak firing. For comparison across environments location within each repeated segment has been converted to a phase (radians). Note the sequence of firing within each environment with no strong preference for any particular phase. **C**, Ratemaps from environments A (*Left*) and B* (*Middle*) sorted according to the order of their peaks in A. (*Right*) Cross-correlation between A and B* shows a significant peak (vs 1000 shuffles, purple line also indicates shuffle confidence interval), suggesting a tendency for the relative phase of ratemaps to be preserved between environment A and B.

## Discussion

The core finding presented here is the report of a distinct population of non-grid neurons in rodent mEC characterised by robust modulation of their firing rate by visual cues presented in linear VR environments – ‘cue-locked’ cells. Indeed, despite the fact that the visual features differed substantially, the majority of neurons not only exhibited strong cue locking in multiple environments but also showed a marked tendency to preserve the relative sequence of their firing fields.

The observation of these cells leads to two main considerations, one with respect to how experiments are conducted in VR and one on the nature of these cells. As previously mentioned, VR has become an increasingly popular tool used to study spatial cognition and its neural basis (Chen et al., 2013; Dombeck et al., 2010; Domnisoru et al., 2013; Harvey et al., 2009; Campbell et al., 2018; Thurley and Ayaz, 2017). In particular, several studies of head-fixed mice on VR linear tracks have considered mEC neurons with multiple similarly sized firing fields to be analogous to grid cells (Campbell et al., 2018; Domnisoru et al., 2013; Schmidt-Hieber and Häusser, 2014). Clearly in linear VR environments without regularly repeating elements cue-locked cells would be expected to generate fewer repetitive fields. Nevertheless, our observation that these cells retain their firing characteristics across environments suggests they would be expected to form multiple fields under many conditions - leading them to be identified as grid cells. Importantly, these findings do not contradict conclusions drawn from previous VR grid cell studies. Indeed, many publications using 1D VR grid cell recordings relied on 2D RW environments for grid cell classification (Campbell et al., 2018; Domnisoru et al., 2013). Moreover, although the 1D VR environments used by Domnisoru et al., 2013 incorporated regularly repeating cues these were constrained to sub-sections of the track. Under such circumstances it is unlikely that cue-locking cells would be confounded with grid cells. Never-the-less, it still appears that the most reliable means of detecting grid cells is via their characteristic six-fold symmetric firing pattern visualised in a 2D environment either real or virtual (Chen et al., 2018; Hafting et al., 2005), though even then the expected false positive error rate is non-negligible (Barry and Burgess, 2017).

In this study, VR environments consisted of long linear tracks (10m) composed of repetitive cues distributed at differing frequencies (67, 100 and 133 cm). As a consequence, the spatial activity of cue-locked cells was remarkably regular (**Figure 3**, **Supp. Fig 4-5**), allowing them to be identified by the strength of their periodicity. Conversely, although we observed that mEC grid cells – identified from open field trials – exhibited multiple firing fields in VR, we found no evidence for cue-locking, corroborating the widely held view that grid cells are strongly modulated by self-motion information (Burgess, 2008; Campbell et al., 2018; Carpenter et al., 2015; Hafting et al., 2005; Winter et al., 2015). In contrast, we found a population of cells showing stable regular firing pattern during VR navigation in register with the available spatial cues. Superficially the visual pattern of the simplest environment (A) resembles drifting sine gratings. However, sensory-motor feedback between the animal’s movement and the visual scene coupled with the spatial perspective and optic flow provided by other textures, suggests that the cells were responding in the context of VR navigation. In light of these considerations, we propose that the striking regular firing pattern exhibited by these cells was predominantly related to visual cues, and hence we consider them to be “cue-locked”. We note that these cells’ spatial fields were not limited to the immediate proximity of cues but spread across the range of possible phases within each repeated environmental segment (**Figure 4**, **Supp. Fig 4-5**) in a way that resembled the sequential firing of an ensemble of grid cells from the same module (Barry et al., 2007; Stensola et al., 2012; Yoon et al., 2016). Interestingly, our results suggest that such sequential firing pattern was conserved at least between two VR environments (A and B, **Figure 4**), suggesting that cue-locking cells may encode relative distances from the cues.

What might be the identity of the cue-locked cells? One possible interpretation is that these neurons are boundary vector cells (BVCs) (Barry et al., 2006; Hartley et al., 2000; Lever et al., 2009) or border cells (Solstad et al., 2008) responding to visual cues that are perceived as a boundary. However, the lack of clear spatial modulation in the open field trials renders this unlikely as BVC firing fields in the open field are typically expected to be unitary and elongated – a simple function of the animals allocentric location relative to environmental boundaries (Hartley et al., 2000; Lever et al., 2009). Without further evidence it is hard to draw solid conclusions. Still it seems plausible that these cells respond to visual features of intermediate complexity and may likely be modulated by the egocentric location of the cue relative to the animal. Although open field recording enclosures often include only a small number of controlled cues like a cue card, it is likely they also include a large number of uncontrolled cues which are unintentionally present. Due to their nature it is necessarily hard to quantify the prevalence and efficacy of such uncontrolled cues. These considerations would account for their relatively simple firing characteristics in the VR in which sensory information is well controlled and behavioral confounds are reduced in contrast to a more complex activity during open field foraging.

Therefore, it falls to future work to further characterise the activity of these cells and the factors they respond to. On one hand it is obvious that the highly simplified and well controlled environments provided by VR have a role in this endeavour but equally less constrained open field recordings combined with careful behavioural tracking will also be important.

## Supporting information

## Acknowledgements

This work was supported by the Wellcome Trust and Royal Society.

