## Supplementary material for "Entorhinal neurons exhibit cue locking in rodent VR"

### Supplementary figures

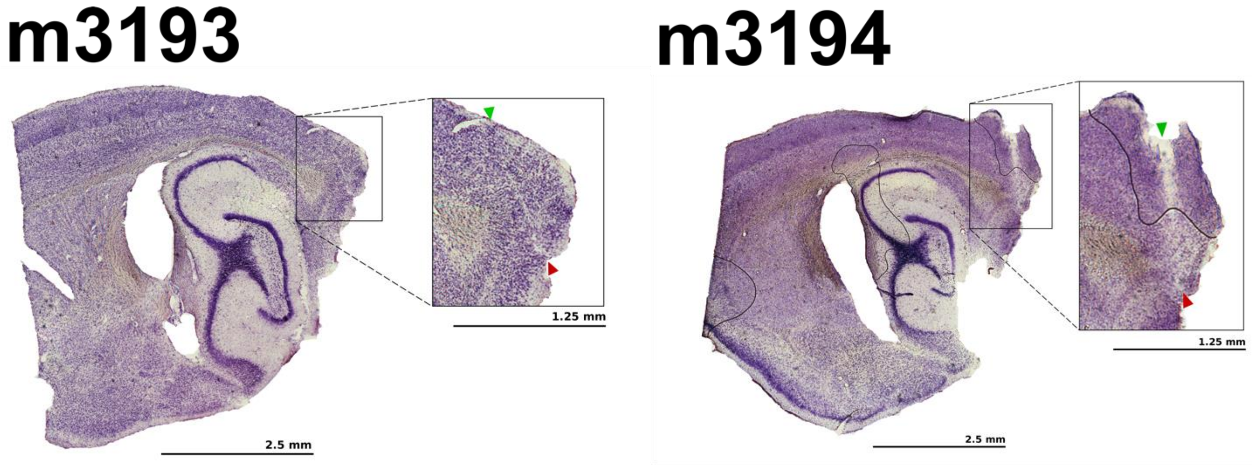


**Supp. Fig 1**

Example histology, showing superficial mEC recording locations in mice m3193 and m3194.

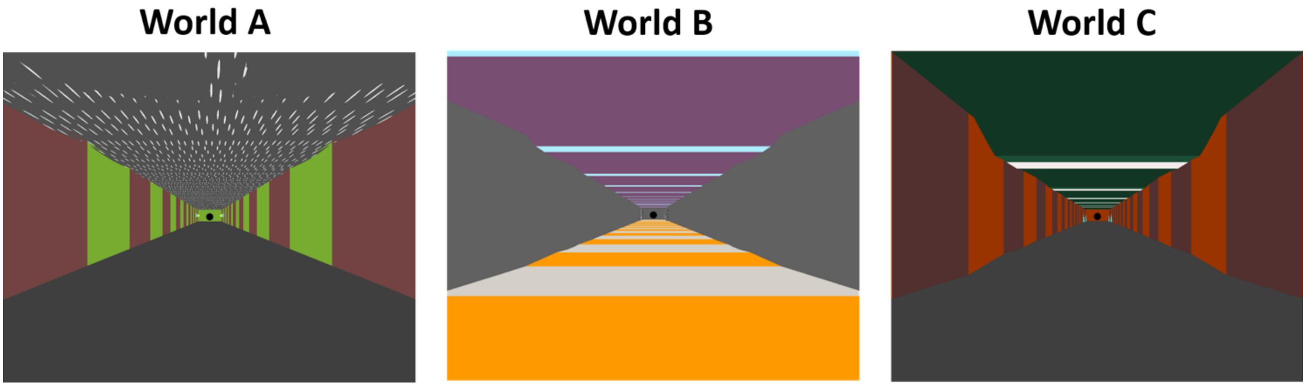


**Supp. Fig 2**

First person perspective view of VR environments A, B, C. Note the differences in the position, colour, and width of the repetitive visual cues across VR environments.

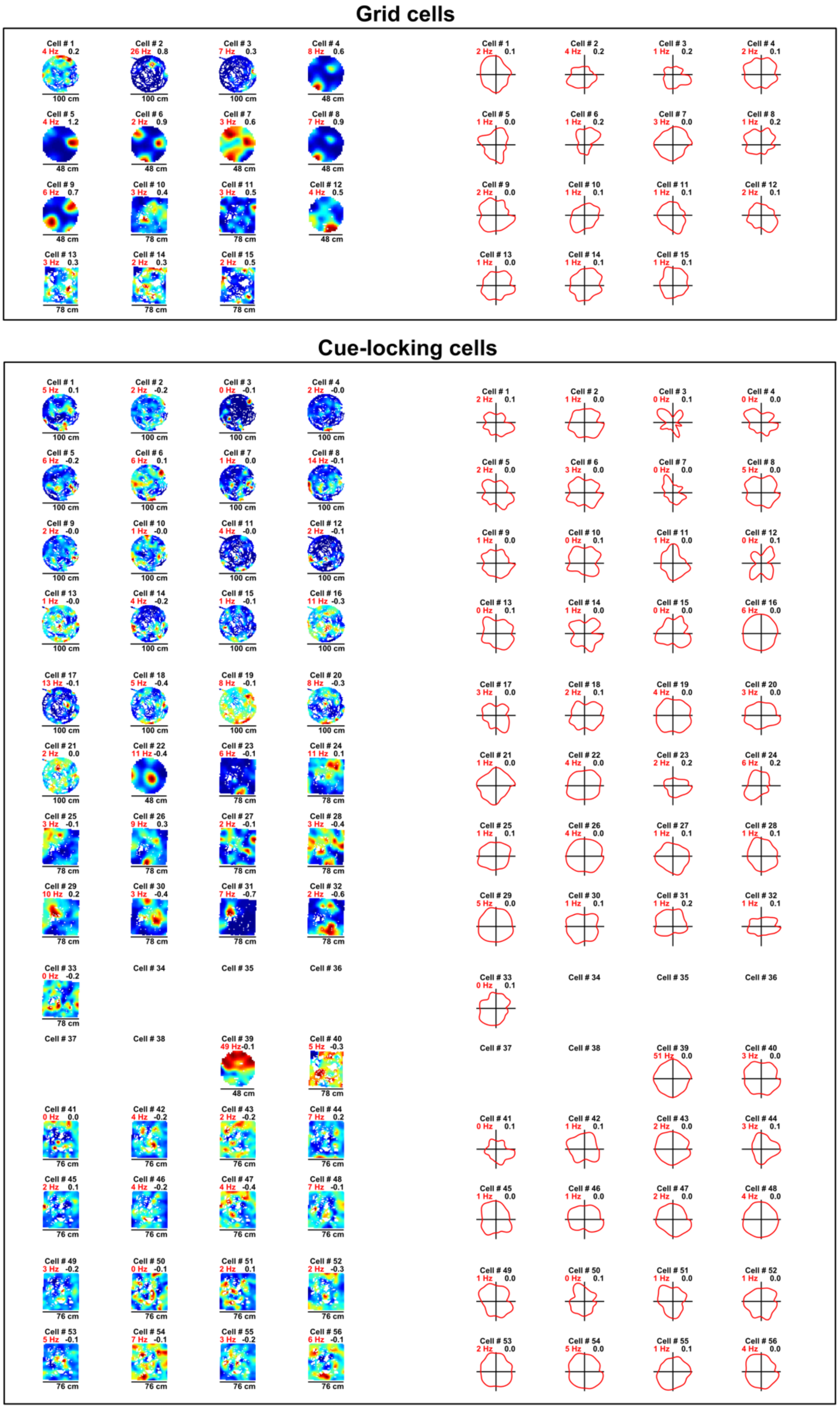


**Supp. Fig 3**

Spatial ratemaps and polar plots of grid cells (*top*) and cue-locking cells (*bottom*) recorded in the open field with peak firing rate of both rate maps and polar plots represented in the top left corner in red. Grid scores and Rayleigh vector length obtained from the corresponding ratemaps and polar plots are shown in the top right corner of each cell.

**Supp. Fig 4**

Rate maps (red lines) of 16 representative examples of periodic cells recorded across VR environments A-C (this page and the one following).

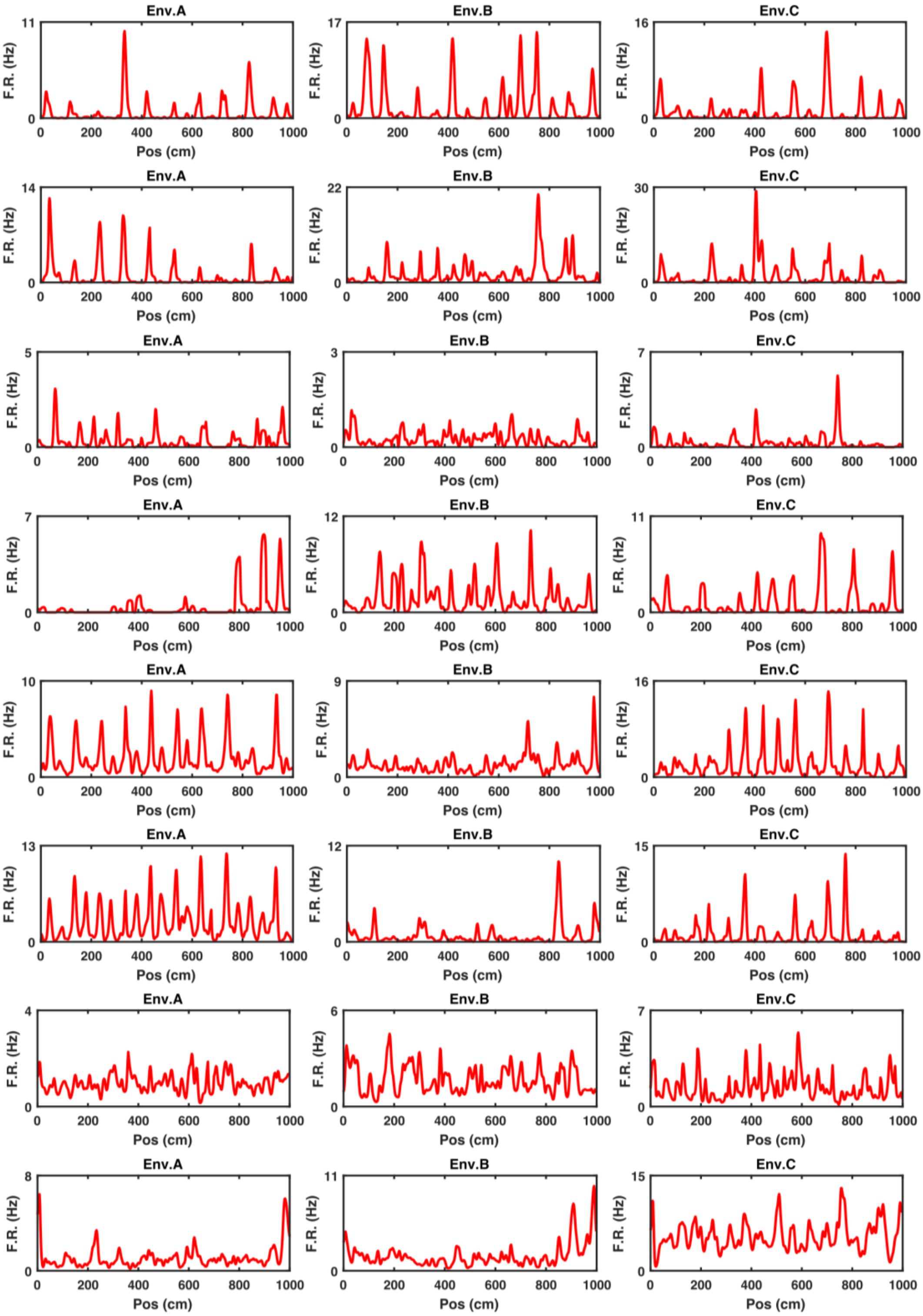

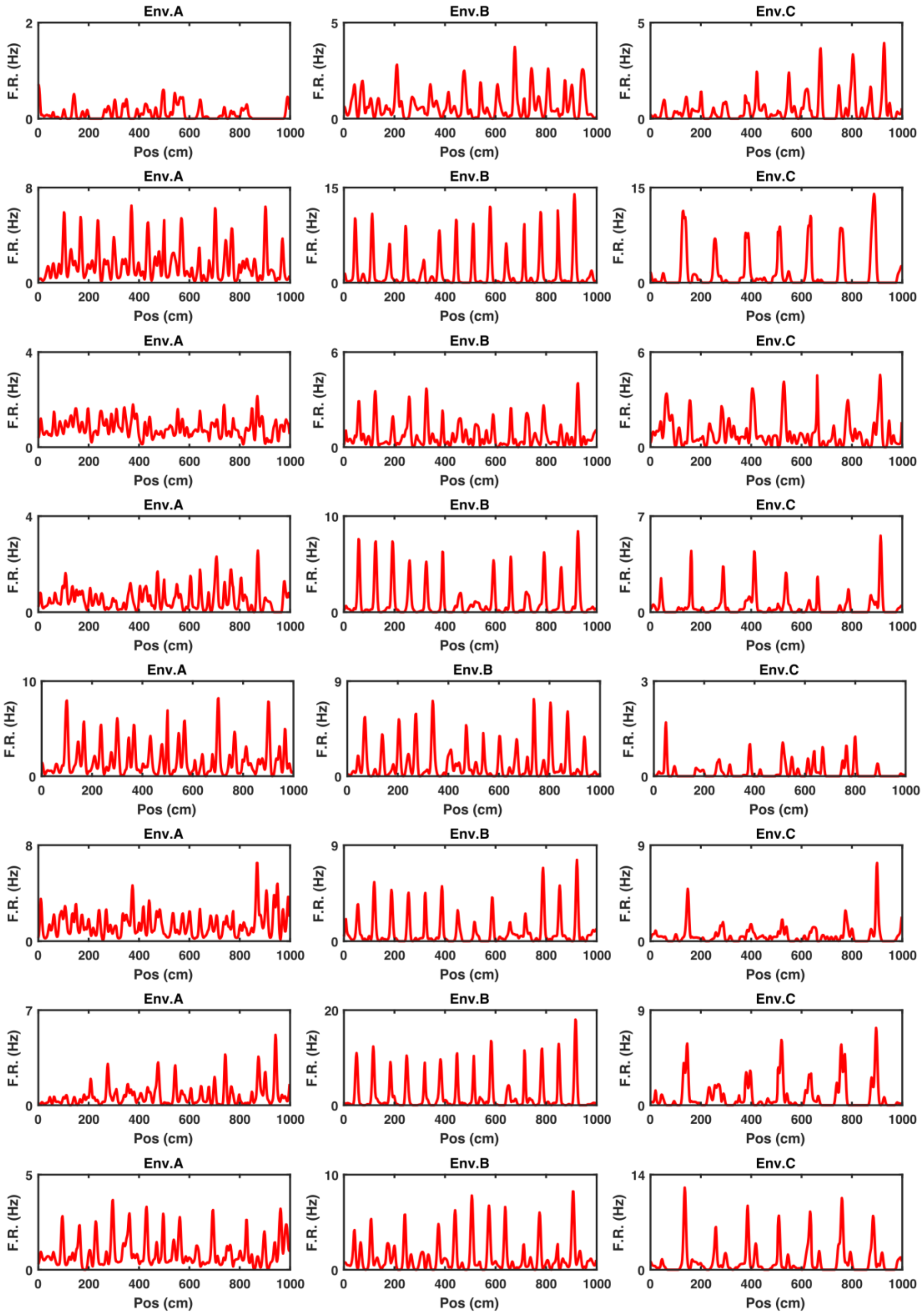


24

**Supp. Fig 5**

Spatial tuning curves (blue lines) of the same 16 representative examples of periodic cells shown in **Supp. Fig 4** calculated as the mean firing rate map over the repeating units for each individual VR environment A-C (this page and the one following).

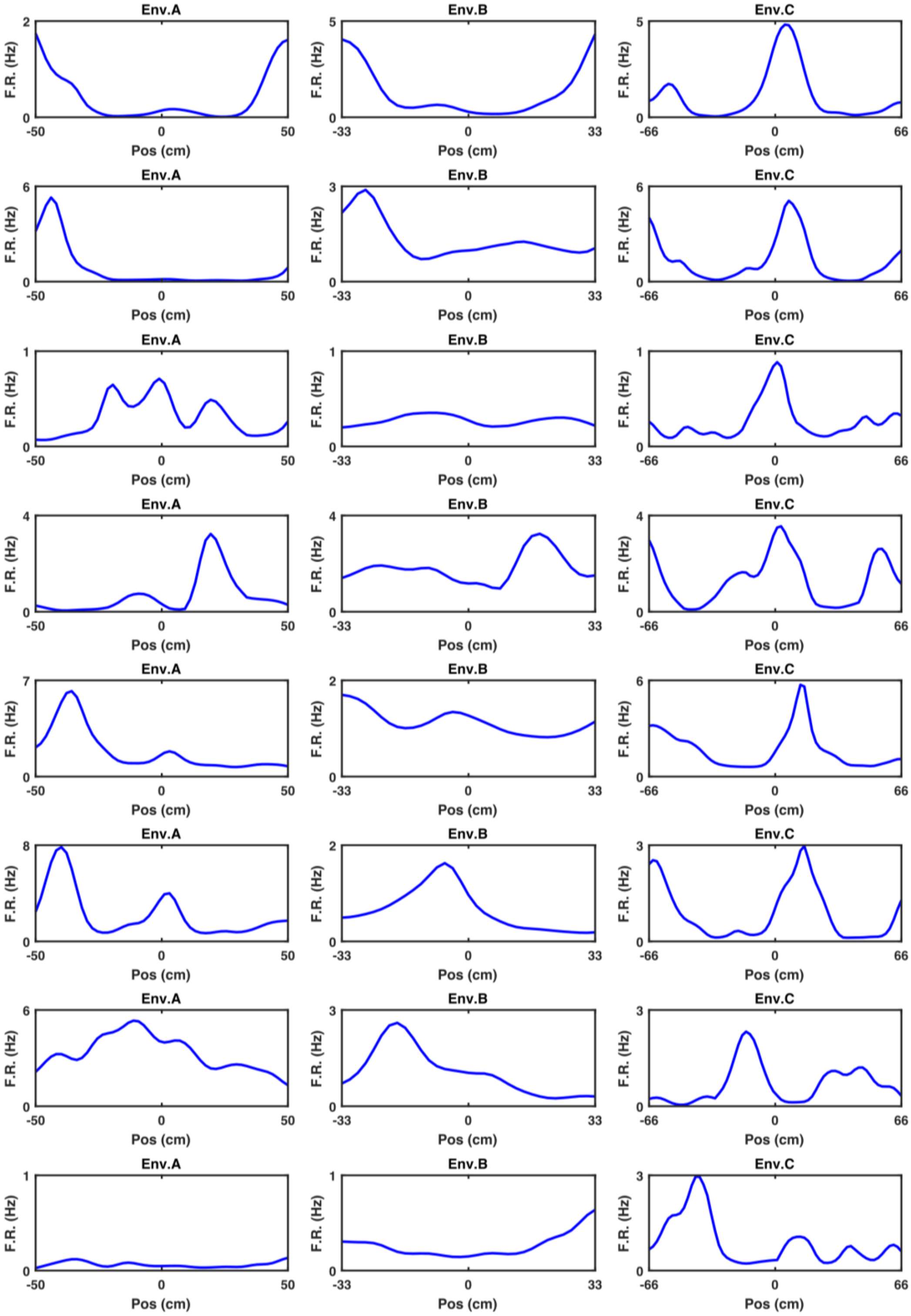

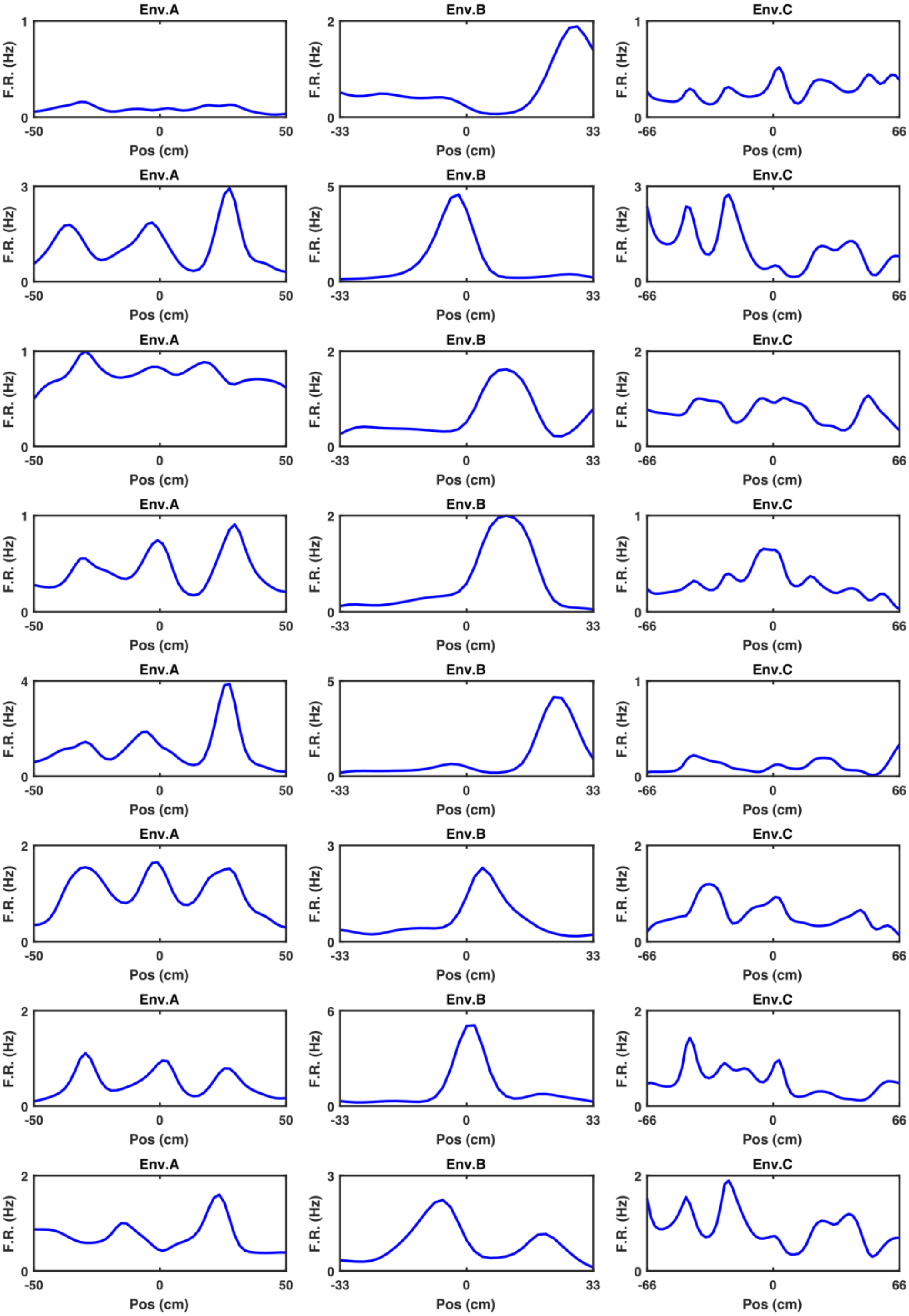

Env.A
Env.B
Env.C
F.R. (Hz)
Pos (cm)
-50
0
50
-33
33
-66
66

### Methods

**Subjects**

Four male C57BL/6J mice aged 7-10 weeks and weighing 25-30 grams at time of surgery were used in this study. Surgeries took place after at least 1 week of habituation in the holding room. All mice were implanted with microdrives housing 8 tetrodes in left MEC (M.L. = 3.2 mm from Lambda, A.P. = 0.4 mm anterior the edge of the sinus). One mouse had additional microdrive implanted in the right HPC (M.L. = 1.8 mm from Bregma, A.P. = 2.0 mm posterior of Bregma). During recovery (1 week from surgery) animals had ad libitum access to food and water and meloxicam (Metacam dogs dosage, 0.1 ml) dissolved in soya milk. Experimentation commenced 7-10 days since surgical operation and coincided with food deprivation regime during which body weight was daily monitored. A minimum weight was set as the 90% of the original weight and increased by 0.5 grams per week since start of food deprivation. During food deprivation, *ad libitum* access to water was maintained. All procedures were licensed by the UK Home Office following the revised ASPA regulations (2013) modified by the European Directive 2010/63/EU.

**Microdrives and single-unit recording**

Extracellular recordings were conducted using Axona (Ltd, UK) microdrives hosting bundles of 8 tetrodes, each composed of four twisted 17μm polyimide-coated platinum-iridium (90%/10%) wires (California Fine Wire, CA) and attached to two 16-channel omnetics connectors compatible with 32-channel Axona headstages (Axona, Ltd,UK).

On recording days, each mouse was brought into the screening room and connected to the recording apparatus (Axona, Ltd, UK) via a litz-wire cable connecting the headstage to the preamplifier. Local field potential (LFP) activity was recorded from 1-8 electrodes, sampled at 250 Hz and bandpass filtered between 0 and 475 Hz. To detect single-unit activity, DacqUSB software (Axona, Ltd, UK) was used in the spike-filtered mode. The signal recorded from each electrode was amplified 8000-38000 times, bandpass filtered between 500 Hz and 7 kHz and sampled at 50 kHz. Spikes were recorded if the voltage signal passed an arbitrary threshold set individually for each individual electrode by the experimenter and it was fixed across RW and VR sessions. During RW navigation, the animal instantaneous position was detected using an infra-red camera (sampling rate = 50 Hz) tracking an array of small LEDs placed on the head-stage and synchronized with the neural signal.

**Experimental apparatus**

VR

Briefly, the VR apparatus consisted of a Styrofoam wheel with head fixation post, lick detector with custom-made reward tube and 4 Iiyama ProLife E1980SD LCD monitors - one placed directly in front of the animals, one on either side and one directly above (**Supp. Fig 6A**). This configuration provides a theoretical field of view spanning 270° horizontally and 180° vertically, comfortably exceeding that used in a number of other studies which have investigated spatially modulated activity in the entorhinal-hippocampal system (Chen et al., 2013; Saleem et al., 2013). The rear was bounded by a curtain.

VR environments were constructed and run on MATLAB 2015a using the ViRMEN (https://pni.princeton.edu/pni-software-tools/virmen) software package. Animal position relative to VR environment was updated online via rotary encoder (Kubler Incremental Encoder 1024 ppr 12000rpm; 160 kHz switching frequency) detecting instantaneous angular velocity of the running wheel. To encourage navigation, milk (sweetened SMA soya formula milk) reward was delivered by a D220S-V micropump (http://micropumps.co.uk) attached to silicone tubing positioned directly in front of the mouse. Reward was delivered via a pinch valve (NResearch Inc) controlled by MATLAB. Licking was detected using an infrared LED (Digi-key, 1080-1357-1-ND) and photodetector (Digi-key, 365-1115-ND) attached on opposite sides of a 3D printed custom-made feeder. Neural activity was recorded using a separate computer running Axona DACQUsb software (Axona, Ltd,UK) and synchronised *post-hoc* by aligning TTL pulses generated in ViRMEN every 10 seconds from the start of the trial.

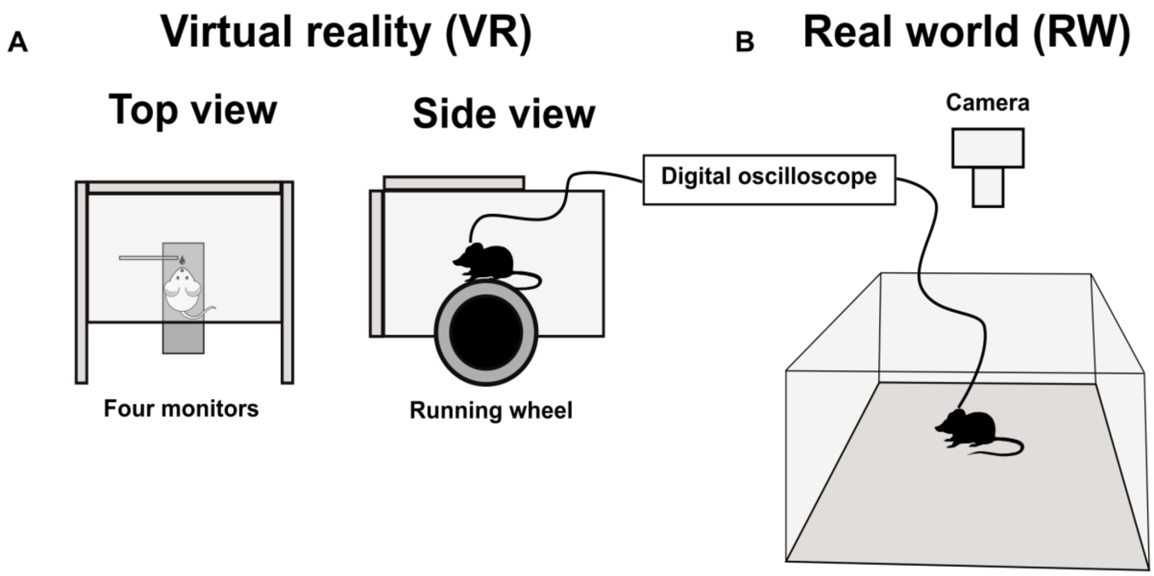


**Supp. Fig 6**

Four-monitor virtual reality apparatus and real world recording setup. (**A**) Schematic representation of the VR apparatus consisting of four adjacent monitors (three horizontal placed front, left and right and one above the animal). Together this configuration provides a field of view spanning 270° horizontally and 180° vertically. A custom-made optical switch placed in front of the animal's snout was used to detect licks. Electrophysiological recordings were conducted by connecting the microdrive implanted on the animal's head to a digital oscilloscope (Axona, Ltd, UK) and synchronizing movements in VR post-hoc. (**B**) Standard open field recordings in RW were conducted similarly with the 2D position of the animal being recorded by an infrared camera.

Animals were recorded while running on three different VR environments (Environment A, Environment B, Environment C) which consistent of 10-meter tracks but differed in frequency, colour, width and location of the available spatial cues (**Supp. Fig 2**). All environments had a dark grey tunnel at the end of the corridor and a reward zone which was indicated by four cues against the walls at 800 and 950 cm along the track.

Alternating green and brown stripes on both of the side walls provided the only repeating visual cue in environment A. Each stripe was 50 mm wide therefore the pattern repeated every 100 mm. The floor of this environment was a plain dark grey and the ceiling was grey with white spots with no clear repeating pattern. The reward zone at the end of the track was indicated by four spheres in yellow and blue.

Environment B had two repeating patterns, both of which repeated every 66.6 mm. Alternating yellow and grey stripes of equal width formed the floor pattern and the ceiling was purple with a thin blue stripe at 40-53 mm (60-80%). At the corners of the reward zone were black columns with white spots.

Environment C also had two repeating patterns however these repeated at different frequencies. The side walls had alternating light and dark brown stripes of equal width which repeated every 67 cm. The ceiling had a pattern which repeated every 133 cm which consisted of a green background with a white stripe of width 33.3 cm (10-25%). The reward zone was cornered by blue and white striped corners.

**<u>Open Field</u>**

Mice foraged for milk (sweetened SMA soya formula milk) droplets in different circular environments (50-100 cm diameter circular arena) or squared 75 cm environments with walls 50 cm high (**Supp. Fig 6B**). The arena was surrounded by black curtains and had a polarizing 5 cm black stripe cue on the east side of the circular environments while a 23 x 55 cm white cue card hung portrait 160 cm above the West wall. During foraging single unit recordings were obtained using a DacqUSB software (Axona, Ltd,UK). Position was recorded by tracking a head-mounted infrared LED using a camera above the centre of the arena.

**<u>Behavioural procedure</u>**

Mice were screened daily for spatially selective cells in the open field foraging task above. Tetrodes were advanced 65-175 μm if no cells of interest were recorded. Immediately after screening, animals were then head-fixed in the VR setup, left in darkness for around 5-10 minutes, and trained to run in an environment consisting of a long 1D VR track (400 cm) with distinct objects (e.g. tower, cylinder, torus) and different patterns on both the floor and the ceiling. During training, the animals received a milk reward in a marked goal location, they then continued moving to the end of the track, entering a tunnel identical to the ones used in the probe environments (A, B and C) before being teleported back to the start of the track. Animals were trained until they could reliably complete 3 laps, this typically meant that animals received at least 5 training sessions, each of which lasted for around 10-20 minutes, before being exposed to the probe VR environments.

During each experimental session, the VR probe environments (A, B, and C) were presented in a random order. Animals were exposed to each environment for 10-30 minutes with a break of 5-10 minutes between environments, during which animals were allowed to rest in the dark. In each environment, animals were required to run along the track and reach a visually-cued reward location 87.5 cm from the start of the track. Animals then ran on to reach a tunnel at the end of the track and, after a random delay ($\sigma = 5$ seconds), were 'teleported' back to the start of the track ready to complete another lap.

### Data analysis

All analyses were conducted on MATLAB 2015a (Mathworks, MA) unless otherwise stated.

**Cluster cutting**

Cluster cutting was performed blindly once for each session by combining multiple sessions in VR with the related open field recordings. This was done to ensure that the neural activity of the same cell could be monitored across multiple recordings without entering any experimenter-based bias in the cell-identfication. Preliminary clusters were assigned via KlustaKwick algorithm (Kadir et al., 2014) across all sessions and then manually corrected by eye based upon spike amplitude, temporal autocorrelogram and waveform of the clusters using Tint software (Axona, Ltd, UK). Once cluster-identification was completed, spikes were then assigned to each recording session and the cut file generated for each individual session in Matlab with custom-made algorithm.

**Spatial analysis**

Spatial rate map

The spatial firing of each individual cell was firstly assessed by examining the spatial rate map (2 cm bin). For each cell, the spike map (overall number of spikes across sampled portion of the environment) and the dwell map (overall dwell time across sampled portion of the environment) were obtained and smoothed using a 5 bin Gaussian smoothing kernel ($\sigma$ = 4 cm). Smoothed spatial rate map representing mean firing rate across space was then obtained dividing the smoothed spike map by the smoothed dwell map. This was done for data in both VR and RW.

Polar rate map

Analogous to the spatial rate map, the directional firing of each individual cell was assessed by producing a polar map (6° bin) obtained by diving a spike map (overall number of spikes across heading direction) by the dwell map (overall dwell time across heading direction). This was done only for data in the RW.

Spatial stability

To determine spatial stability of the ratemaps in both VR and RW within the session, for each cell the spatial ratemaps of the first and second half of the trial duration were generated. The stability was quantified as the *Pearson* correlation between corresponding bins with firing rate > 0 Hz of the fist vs second half of the session.

Spatial information

The spatial content of each cell was determined using the Skaggs spatial information index (Skaggs et al., 1993) and obtained as follows:

$$I(R|X) \approx \sum_i p(x_i)\ f(x_i)\ log_2\left(\frac{f(x_i)}{F}\right)$$

where $p(x_i)$ and $f(x_i)$ correspond to the probability of the animal being in location (spatial bin) $x_i$ and the firing rate of cell in location $x_i$, respectively. *F* is the overall firing rate and $I(R|X)$ corresponds to the amount of spatial information between firing rate *R* and location *X.* The spatial information in bits/spike was calculated by dividing the $I(R|X)$ by *F*.

Spatial autocorrelogram (SAC) in RW

To determine the periodicity of the spatial firing of each individual cell, the SAC was conducted using the following formula (Hafting et al., 2005):

$$r(\tau_x, \tau_y) = \frac{n\Sigma\lambda(x,y)\lambda(x-\tau_x, y-\tau_y) - \Sigma\lambda(x,y)\lambda(x-\tau_x, y-\tau_y)}{\sqrt{n\,\Sigma\lambda(x,y)^2 - (\Sigma\lambda(x,y))^2}\sqrt{n\Sigma\lambda(x-\tau_x, y-\tau_y)^2 - (\Sigma\lambda(x-\tau_x, y-\tau_y))^2}}$$

where $r(\tau_x, \tau_y)$ is the correlation coefficient between those bins with spatial offset $\tau_x$ and $\tau_y$, $\lambda(x, y)$ is the firing rate in the spatial bin of the spatial rate map defined by coordinate $x$ and $y$, and $n$ is the total number of spatial bins. The grid score for RW data was obtained after detecting the 6 nearest peaks from the centre of the SAC and subtracting the lowest correlation found at 60° and 120° to the maximum correlation at 30°, 90° and 150° relative to the orientation of the first peak.

**Spatial frequency**

To determine the spatial frequency in VR, a spatial rate map of each trial was produced and used to generate a linear SAC of a single trial in the 20-180 cm range. This process was repeated for every trial completed by the animal in each environment so that a mean SAC across trials could be obtained by averaging SAC of each individual trial.

To characterize the putative periodic firing of a cell, the mean SAC was analysed using the inbuilt function *findpeaks* in Matlab 2015. The strength and spatial frequency of each cell in VR was determined respectively as the “prominence” of the mean SAC highest peak and the distance at which the mean SAC showed highest peak. A shuffling procedure was conducted by shifting the cell spike-train by a random delay > 30 seconds. Shuffled rate maps of each trial were then generated and used to produce a mean shuffled SAC, so that a shuffle prominence value could be obtained. This process was repeated 1000 times for each cell.

<u>Cell identification criteria</u>

A cell passed periodic cell acceptance criteria if all the following criteria were met on at least 2/3 VR environments: mean firing rate < 5 Hz, peak firing rate > 1.5 Hz, prominence of the highest peak detected from the mean SAC exceeding the 95perctile of the shuffled distribution (*n* shuffles = 1000).

A cell passed grid cell acceptance criteria if the following criteria were met in the RW open field: mean firing rate < 3 Hz, peak firing rate > 1 Hz, spatial stability > 0.2, grid score exceeding 95perctile of the shuffled distribution (*n* shuffles = 200) obtained by shifting the spike-train relative to dwell time by a random amount > 30 seconds.
